# Global distribution and diversity of marine Parmales

**DOI:** 10.1101/2023.11.09.566328

**Authors:** Hiroki Ban, Hisashi Endo, The EukBank Team, Akira Kuwata, Hiroyuki Ogata

## Abstract

Parmales (Bolidophyceae) is a minor eukaryotic phytoplankton group, sister to diatoms, that exists as two distinct forms of unicellular organisms: silicified cells and naked flagellates. Since their discovery, many field studies of Parmales have been performed, but their global distribution remains under-investigated. Here, we compile over 3,000 marine DNA metabarcoding datasets targeting the V4 region of the 18S rRNA gene from the EukBank database. By linking this large dataset with the latest morphological and genetic information, we provide updated estimates on the diversity and distribution of Parmales in the global ocean at a fine taxonomic resolution. Parmalean amplicon sequence variants (ASVs) were detected in nearly 90% of the analyzed samples. However, the relative abundance of parmaleans in the eukaryotic community is less than 0.2% on average, and the estimated true richness of parmalean ASVs is about 316 ASVs, confirming their low abundance and low diversity. Phylogenetic analysis clearly divides these algae into four clades, and three known morphotypes of silicified cells were classified into three different clades. The abundance of Parmales is generally high in the pole and decreases toward the tropics, and individual clades/subclades show further distinctions in distributions. Overall, our results suggest their clade/subclade-specific adaptation to different ecological niches.

## Introduction

The order Parmales (class Bolidophyceae) comprises eukaryotic microalgae of two morphologically distinct forms: one form is a naked flagellate (1–1.7 μm) and the other form has silicified cell walls (2–5 μm) (Booth and Marchant, 1987; Guillou et al., 1999; Ichinomiya et al., 2011, 2016). The silicified forms were originally established as a new order, Parmales, within Chrysophyceae (Booth and Marchant, 1987). Following the first isolation of a parmalean from the Oyashio region near Japan (Ichinomiya et al., 2011), phylogenetic analyses revealed that parmaleans form a monophyletic group with the previously known naked flagellates (bolidophytes) (Ichinomiya et al., 2011, 2016; Tajima et al., 2016) that comprise a sister group to diatoms (Guillou et al., 1999). Consequently, the order Parmales was re-established under the class Bolidophyceae (Ichinomiya et al., 2016). The two forms have a phylogenetically nested relationship, and the silicified strains possess the genomic potential to form flagella, suggesting that these two forms represent different stages in the life cycle of the same organisms (Ichinomiya et al., 2016; Yamada et al., 2020; Ban et al., 2023). Currently, there are four identified morphotypes of silicified parmaleans, each distinguished by the morphological features of their silicified cell walls: *Triparma*, *Tetraparma*, ‘Scaly parma’, and *Pentalamina*, of which the last morphotype has not yet been isolated (Booth and Marchant, 1987; Ban et al., 2023; Sato et al., Unpublished).

Their life cycle and predicted phago-mixotrophic nutrient acquisition contrast sharply with those of diatoms, their closest evolutionary photo-autotrophic relatives (Ban et al., 2023). Therefore, Parmales is a key eukaryotic group in understanding the physiology, ecology, and evolution of diatoms, the most successful phytoplankton group in the modern ocean (Kuwata et al., 2018; Ban et al., 2023). In particular, characterizing the diversity and biogeography of Parmales across space and niches is expected to provide fundamental information on the difference in the ecological and evolutionary strategies between diatoms and Parmales.

Based on field observations to date, the silicified form of Parmales is widely distributed, from frequently-reported polar and subpolar regions including coastal sites (Booth et al., 1980, 1981; Silver et al., 1980; Nishida, 1986; Taniguchi et al., 1995; Komuro et al., 2005; Konno et al., 2007; Ichinomiya et al., 2010, 2019; Ichinomiya and Kuwata, 2015; Luan et al., 2018) to the tropics (Kosman et al., 1993; Bravo-Sierra and Hernandez-Becerril, 2003; Fujita and Jordan, 2017). However, these observations were based on microscopic analyses of silicified cell wall morphology, and thus naked flagellates were missed in the observations. There were also potential issues regarding cryptic species (Bickford et al., 2007) and variation within a single species due to morphological phenotypic plasticity (Konno et al., 2007). Thus, an accurate and consistent taxon identification method is needed to quantify the abundance and diversity of Parmales.

Recently, DNA metabarcoding targeting the V4 or V9 region of the 18S rRNA gene has become an effective method to explore the eukaryotic diversity and community composition of the ocean (De Vargas et al., 2015; Massana et al., 2015; Cordier et al., 2022). DNA metabarcoding bypasses microscopic taxonomy identification and provides abundant comprehensive data that may resolve the issues with microscopic observation. Previous studies have investigated the distribution of Parmales using the V9 metabarcoding dataset produced by the *Tara* Oceans expedition (Ichinomiya et al., 2016) and the V4 metabarcoding datasets from multiple studies covering coastal, Arctic, and Antarctic oceans not represented in the *Tara* Oceans data (Kuwata et al., 2018). These studies characterize the global distribution of Parmales in the ocean, revealing their consistently low frequency in microeukaryotic communities and suggesting that each clade of Parmales has its own distribution pattern. However, the data used by Ichinomiya et al. (2016) from the *Tara* Oceans expedition do not cover coastal areas and some oceanic regions such as the Indian Ocean and Antarctic Sea, and the data used by Kuwata et al. (2018) underrepresent the Southern Hemisphere. Additionally, when these studies were conducted, the sequence information of isolated strains was only available for one clade, *Triparma*, of the four morphotypes.

In this study, we use the EukBank database, which provides by far the largest dataset for DNA metabarcodes targeting the 18S rRNA gene V4 region (Kaneko et al., 2023), and associate these data with the current knowledge of Parmales, including information from recent isolates from *Triparma*, *Tetraparma*, and ‘Scaly parma’ (Yamada et al., 2020; Ban et al., 2023). The EukBank project consolidates multiple datasets originating from various sampling projects including *Tara* Oceans (Pesant et al., 2015; Sunagawa et al., 2020), Malaspina (Logares et al., 2020), and Australian Microbiome (Brown et al., 2018). The dataset comprises over 15,000 DNA metabarcoding samples targeting the 18S rRNA gene V4 region, with samples derived from various biomes such as marine, freshwater, and soil biomes. After appropriate filtering, we compiled over 3,000 marine samples from pole-to-pole oceanic regions, including coastal areas. Our results quantitatively describe the diversity and distribution of Parmales at fine taxonomic resolution.

## Material and Methods

### EukBank data preprocessing

The EukBank database provides amplicon sequencing variant (ASV) sequences and their taxonomic annotations, and also includes table data representing read counts of ASVs in each sample with metadata. Detailed information on the preprocessing of raw data is provided in Kaneko et al. (2023).

First, we selected seawater samples with metadata and a total read count over 10,000. Subsequently, we filtered the samples based on the availability of sampling site information on latitude, longitude, and depth. During this process, samples without specific depth information were also retained if the range of sampling depth could be determined from other information (Supplementary Information). Depth information was finally classified as either surface layer (0– 10 m) or euphotic zone (10–200 m). We subsequently retained only samples (depth 0–200 m) with a lower limit of size fraction being less than 1 μm. This size threshold was determined with the expectation that it would allow all parmalean morphological forms to be captured on the filter. Next, we categorized the samples based on their seafloor depth at the sampling sites; sites shallower than 200 m were defined as coastal ocean sites, while deeper sites were defined as open ocean sites. The depths of the sampling sites were calculated by interpolating the depth data from the global relief model (ETOP 1) (Amante, 2009) using latitude and longitude information with the Julia package “Interpolations.jl”. Finally, we collected 3,200 marine samples, of which 1,633 were coastal ocean samples and 1,567 were open ocean samples (Fig. S1).

We also obtained 432 ASV sequences that were annotated as “Bolidophyceae” from the total EukBank ASV sequences.

### Phylogenetic analyses and ASVs annotation

We collected full-length parmalean 18S rRNA genes from the SILVA database (categorized as “Bolidomonas”; accessed Jun 2023) (Quast et al., 2012) and from published work (Ban et al., 2023). We added some diatom sequences as outgroups to the dataset and then removed previously reported chimeric sequences (Ichinomiya et al., 2016) and sequences shorter than 900 bp. We aligned and masked the sequences using the “ssu-align” and “ssu-mask” commands of SSU-ALIGN (v 0.1.1) (Nawrocki, 2009) with default parameters. A maximum likelihood tree was estimated with a GTR + G + F model and 1000 bootstrap replicates using RAxML-ng (v 1.0.2) (Kozlov et al., 2019). We defined clades and subclades based on the topology of the estimated phylogenetic tree (Fig. 1).

**Fig. 1.**
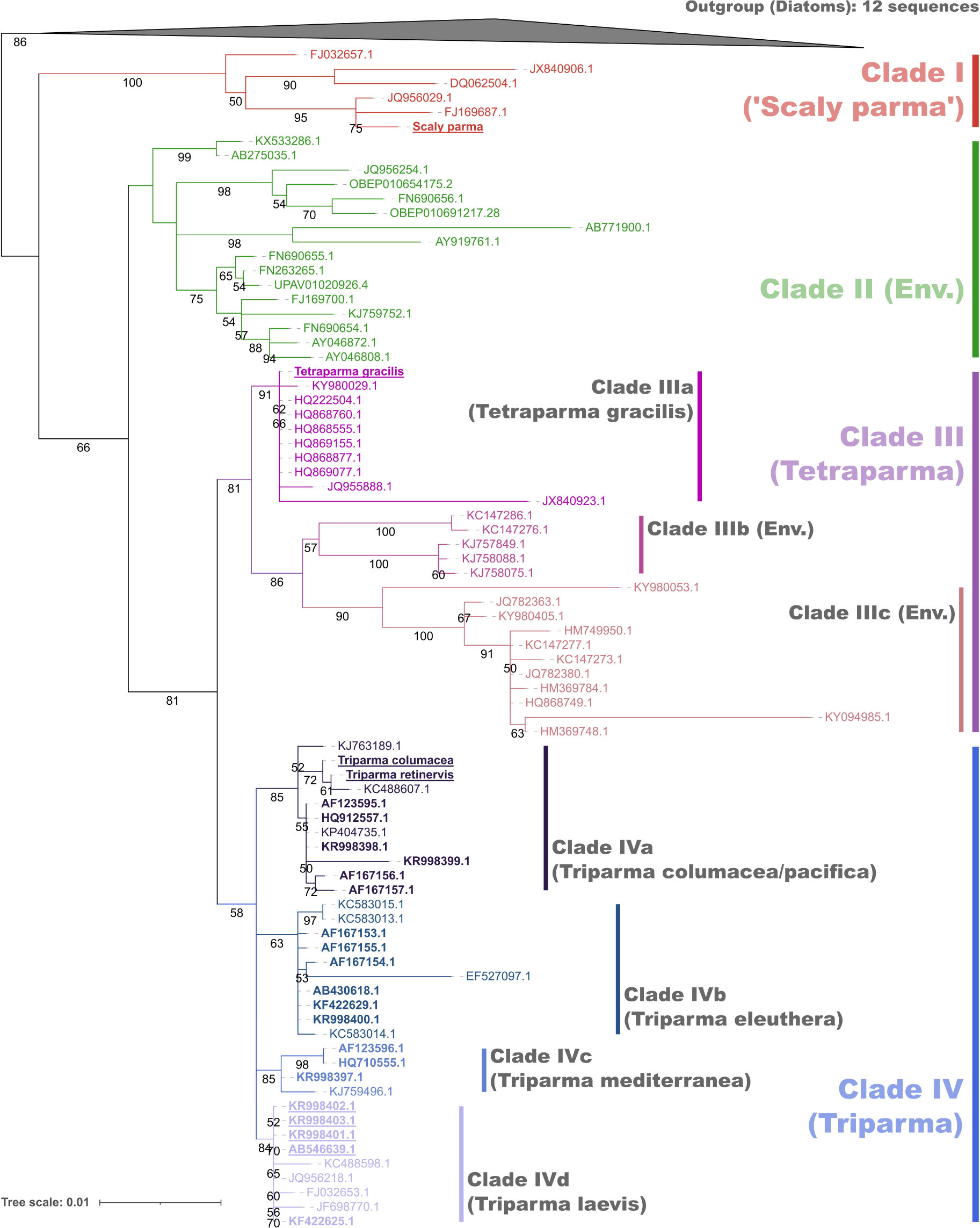
Phylogenetic tree of 18S rRNA genes. Maximum likelihood phylogenetic tree of the full-length 18S rRNA genes of Parmales and diatoms (outgroup). IDs of isolates are in bold and, if isolates are of the silicified form, IDs are underlined. Bootstrap values > 50 are noted on the nodes. The clades/subclades were separated based on the tree topology.

Next, ASVs were assigned phylogenetic placement on the estimated reference tree. First, the full-length 18S rRNA gene sequences were aligned to make a reference multiple sequence alignment using the “ssu-align” command with default parameters, but default masking causes a loss of resolution in distinguish ASVs, so masking of the alignment was done using the “ssu-mask” command with the “--rfonly” option. Sequences of 432 parmalean ASVs were also aligned and masked using the “ssu-align” command with default parameters and the “ssu-mask” command with the previously produced mask file from the full-length 18S rRNA gene alignments. Model parameters for phylogenetic replacement were evaluated with the full-length 18S rRNA gene alignment and the estimated tree using RAxML-ng (Kozlov et al., 2019) with the “--evaluate” option (v 1.0.2) specifying the GTR + G +F model. Phylogenetic placement was done using EPA-ng (Barbera et al., 2019) under the evaluated model with the ASV alignment as the query and the maximum likelihood tree and the full-length 18S rRNA gene alignment as the reference.

To annotate each ASV into a clade/subclade, we used the “extract” command of gappa (v 0.8.4) (Czech et al., 2020). ASVs were annotated into Clades I–IV, “basal_branches”, and “outgroup”. Here we prioritize EukBank’s taxonomy, so ASVs annotated as “outgroup” are treated as Parmales origin, and those annotated as “basal_branches” and “outgroup” are grouped together as “uncertain” sequences of Parmales. At the subclade level, Clade III was divided into Clades IIIa–IIIc, and Clade IV was divided into Clades IVa–IVd (Fig. 1). ASVs annotated as Clade III or IV at the clade level but annotated as “basal_branches” at the subclade level were re-annotated as “Clade III uncertain” or “Clade IV uncertain”, respectively.

The parmalean ASV sequences were also aligned to the full-length 18S rRNA gene sequences using vsearch (v 2.22.1) (Rognes et al., 2016).

### Ecological analyses

The rarefaction curves were obtained by plotting the expected count of ASVs calculated under the assumption that all reads from 3,200 samples were pooled and then sub-sampled. The number of reads sub-sampled increased in increments of 10,000. The slopes of the rarefaction curves were calculated from the last data points and its predecessors.

Preston’s log-normal distribution (Preston, 1948) was used to estimate the completeness of sampling by fitting a left-truncated normal distribution to the log2-transformed total counts of each ASV in 3,200 samples using function “prestondistr” in the R package “vegan” via Julia package “RCall.jl”.

The relationships between latitude and parmalean ASV abundances were visualized by LOESS (Locally Estimated Scatterplot Smoothing; span = 0.8) using the Julia package “Loess.jl”. Confidence intervals were computed using 100 bootstrap resampling iterations.

To characterize the habitats of each clade and subclade, the weighted average temperature (WAT) was used as the index for clade/subclade temperature preferences using the relative abundance in the 2002 of 3200 samples for which temperature information was available. This index was calculated using the following equation:

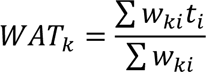

where *WAT_k_* is the weighted average temperature index of clade/subclade *k*, and *w_ki_* and *t_i_* are the relative abundance of clade/subclade *k* and water temperature in sample *i*, respectively. Plots were generated using the Julia packages “Makie.jl” (Danisch and Krumbiegel, 2021) and “GeoMakie.jl”.

### Statistical test

For each clade and subclade, the Mann-Whitney U test (Mann and Whitney, 1947) was employed using the Julia package “HypothesisTests.jl” to evaluate whether a difference exists in relative abundance between coastal ocean samples (n=1633) and open ocean samples (n=1567). The results were further validated using the rank-biserial correlations (RBC), which is an effect size of the Mann-Whitney U test (Wendt, 1972; Kerby, 2014). Positive RBC values indicate a preference for the coastal ocean and negative values indicate a preference for the open ocean; the absolute value (0–1) indicates the strength of the preference.

## Results

### Phylogenetic analyses

The maximum likelihood tree (Fig. 1) based on the phylogenetic analysis of the full-length 18S rRNA gene sequences showed clear grouping of parmalean sequences into four clades (Clade I, Clade II, Clade III, Clade IV), which is consistent with previous findings (Kuwata et al., 2018). Clade I is the most basal clade, containing “Scaly parma” that are morphologically distinct from other parmalean taxon (Ban et al., 2023; Sato et al., Unpublished). Clade II consists only of environmental sequences and does not include any sequences from isolated or cultured strains of either silicified forms or naked flagellates. Clade III was divided into three subclades based on topology (Clade IIIa, Clade IIIb, Clade IIIc). Clade IIIa contains a silicified isolate sequence (*Tetraparma gracilis*) and a sequence of unknown morphology (KY980029: *Triparma pacifica* isolate NY13S_157), and Clade IIIc contains two sequences of unknown morphology (KY980053: *Triparma pacifica* isolate NY13S_197; KY98405: *Triparma pacifica* isolate BH65_151). Clade IIIb consists only of environmental sequences. The taxonomic annotation of *Triparma* for the three sequences of unknown morphology in Clade III may be misannotation of samples that properly belong to *Tetraparma*. Clade IV was also divided into four subclades (Clade IVa, Clade IVb, Clade IVc, Clade IVd). Clade IVa and Clade IVd contain sequences from silicified isolates and naked flagellates, with *Triparma columacea* and *Triparma retinervis* representing the silicified form and *Triparma pacifica* RCC205 (HQ912557) representing the naked flagellate for Clade IVa. In Clade IVd, *Triparma laevis* f. *inornata* (AB546639) represents the silicified form and *Triparma sp.* RCC1657 (KF422625) represents the naked flagellate. Clade IVb and Clade IVc do not have any sequences from silicified form isolates, with *Triparma eleuthera* (KR998400) and *Triparma mediterranea* (KR998397) representing naked flagellates for Clade IVb and Clade IVc, respectively.

As a result of phylogenetic placement of ASVs in the reference maximum likelihood tree, 86.3% of ASVs were annotated into one of the four clades (Table 1). There were no ASVs that matched best with outgroup sequences in the sequence similarity search, suggesting that all ASVs are derived from Parmales. The ASVs that were annotated into clades, as well as those that were not annotated, had sequence identities of 91.7% and 83.1%, respectively, with their closest matching parmalean sequences (Fig. S2). Clade II, the environmental clade, contained the most annotated ASVs, followed by Clade I, Clade III, and Clade IV (Table 1).

**Table 1.** Counts of parmalean ASVs in each annotation category.

| **Annotation** | **No. of parmalean ASVs in all EukBank samples** | **No. of parmalean ASVs in the 3200 marine samples** |
| --- | --- | --- |
| Clade I | 76 | 32 |
| Clade II | 214 | 120 |
| Clade III | 50 | 44 |
| Clade IV | 33 | 28 |
| Uncertain | 59 | 43 |
| **Total** | **432** | **267** |

### Global marine dataset of parmalean ASVs

After filtering samples from the EukBank database, we finally obtained a total of 3200 marine water samples containing about 402 million reads. Although the coastal ocean samples were predominantly from Europe and Australia, the open ocean samples range widely from pole to pole, with some exceptions (e.g., the West Pacific) (Fig. S1). From 432 EukBank ASVs, 267 ASVs appeared at least once and accounted for about 630,000 of the total reads. Overall, 94.6% of parmalean ASV reads were assigned to four clades (Fig. 2a), of which Clade IV was the most dominant, followed by Clade III, Clade II, and Clade I (Fig. 2a). This order of abundance is not consistent with the order of the diversity inside the clades (Table 1). Within Clade III, Clade IIIc was slightly more abundant than Clade IIIa, with Clade IIIb being the least abundant (Fig. 2a). Within Clade IV, Clade IVa was the most abundant, followed by Clade IVd and Clade IVb, with Clade IVc being substantially less abundant (Fig. 2a). Rarefaction analysis indicated that the ASV richness of all global ocean parmaleans was far from saturation (Fig. 2b). Particularly, Clade II had the largest slope (Table S1). The fitted Preston model (blue line in Fig. 2c) extrapolated the true parmalean ASV richness to 315.85 ASVs, indicating that the analyzed samples uncover ∼ 84.5% of parmalean ASVs in the global ocean (right side of the veil line in Fig. 2c).

**Fig. 2.**
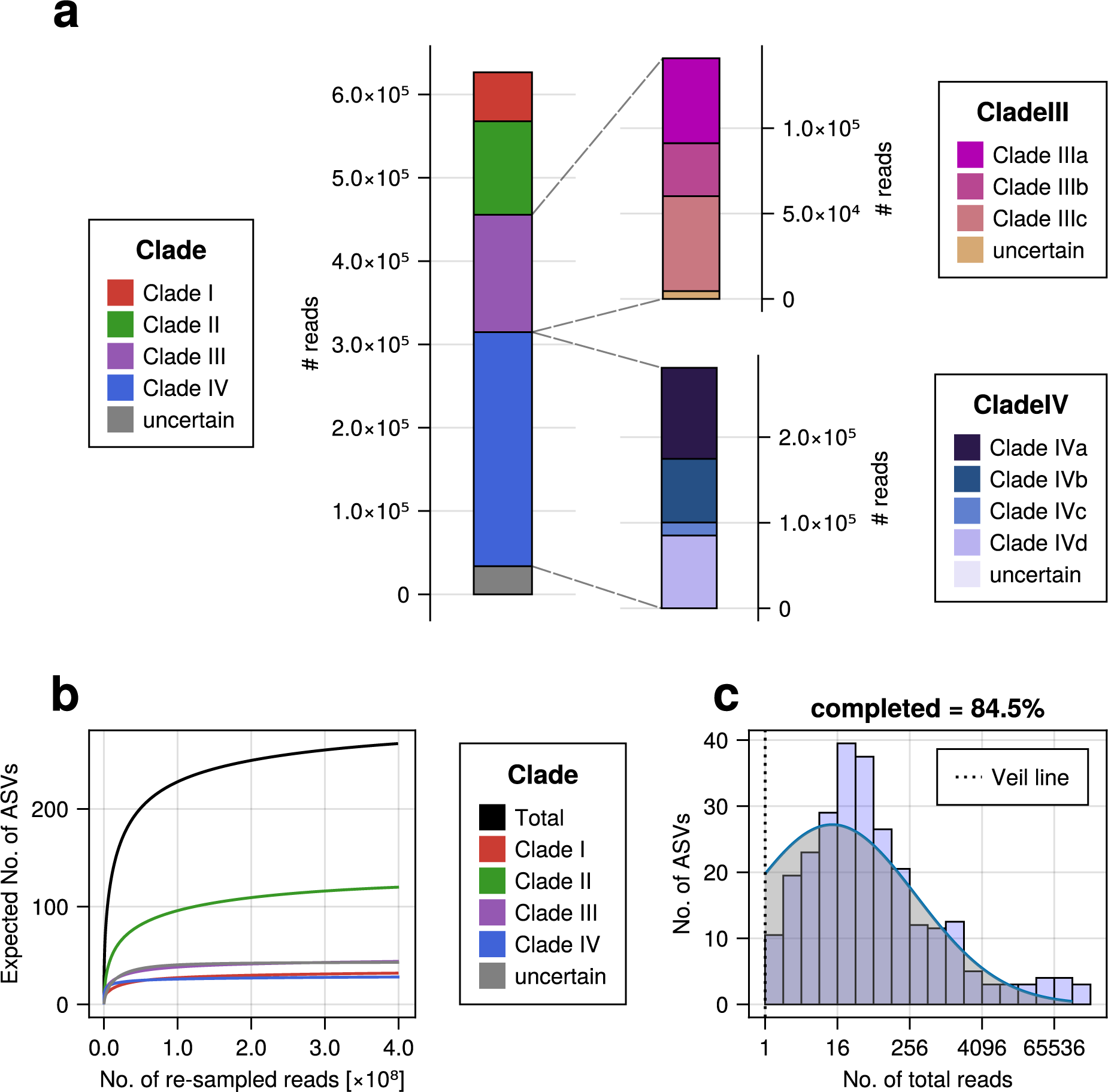
Overview of the parmalean ASV dataset. (a) The number of reads of parmalean ASVs in each clade from 3200 samples. (Top right) The number of reads of parmalean ASVs in Clade III. (Bottom right) The number of reads of parmalean ASVs in Clade IV. (b) Rarefaction curve, representing parmalean ASV richness. Each curve shows the expected number of parmalean ASVs against the number of re-sampled reads for each clade. The slopes of each curve are listed in Table S1. (c) Preston’s log-normal distribution of parmalean ASVs. The x-axis is transformed to log2. The histogram represents the frequency of actual parmalean ASVs binned by abundance in octaves. The blue line indicates the fitted left-truncated normal distribution. The left side of Preston’s Veil line (dashed black vertical line) corresponds to ASVs that did not appear in the samples. The total parmalean ASV richness was extrapolated to ASVs on this side.

### Oceanic distribution of parmalean ASVs

Parmalean ASVs were distributed widely across the global ocean in both coastal and open oceans from the poles to the tropics (Fig. 3a). Parmalean ASVs appeared in 89.1% of the samples, indicating their wide distribution. However, the relative abundance of parmalean ASVs in the microeukaryotic community was generally low, with a median of 0.05% and an average of 0.16% (Fig. 3b). There were exceptional samples where parmalean ASVs were a large portion of the community (three outliers in Fig. 3b). In three samples taken at the same time and location in Botany Bay of Sydney, Australia, parmalean ASVs contributed 58.2%, 48.1%, and 36.5% of the eukaryotic community (SRA Run: SRR8820967, SRR8820804, SRR8820815 from Australian Microbiome). ASVs with a high sample coverage (i.e., the proportion of samples in which they were detected) tended to a have high maximum relative abundance (Fig. 3c). Yet, an ASV that was dominant in one sample was not necessarily widely distributed.

**Fig. 3.**
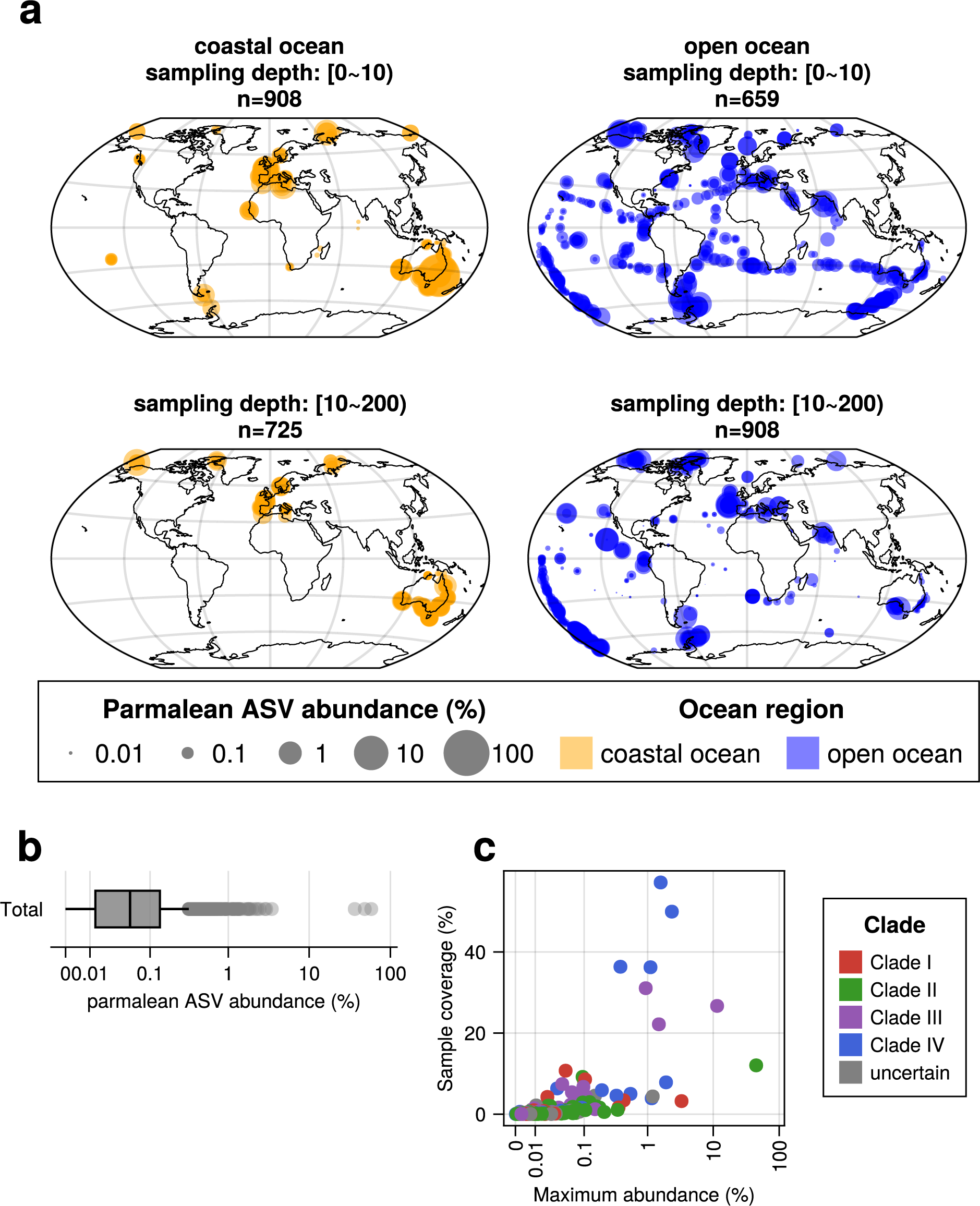
Overview of the parmalean distribution in the global ocean. (a) Global distribution of parmalean ASVs. Top left, Coastal ocean surface layer; Bottom left, Coastal ocean euphotic zone; Top right, Open ocean surface layer; Bottom right, Open ocean euphotic zone. Marker sizes are scaled to the log10(total relative abundance*10,000+1) at each sample. (b) Total relative abundance of parmalean ASVs of each sample. The x-axis is scaled with the pseudolog10 function. (c) Existing ratio and the maximum relative abundance of each ASV. The x-axis is scaled with the pseudolog10 function.

The relative abundance of parmalean ASVs across latitudes showed a clear previously undescribed pattern that decreased from the poles to tropics, while they were detected in the coast and open oceans across the surface and euphotic zones (Fig. 4). In the open ocean, the relative abundance of parmalean ASVs decreased more markedly in the tropics in the euphotic layer than in the surface layer (Fig. 4).

**Fig. 4:**
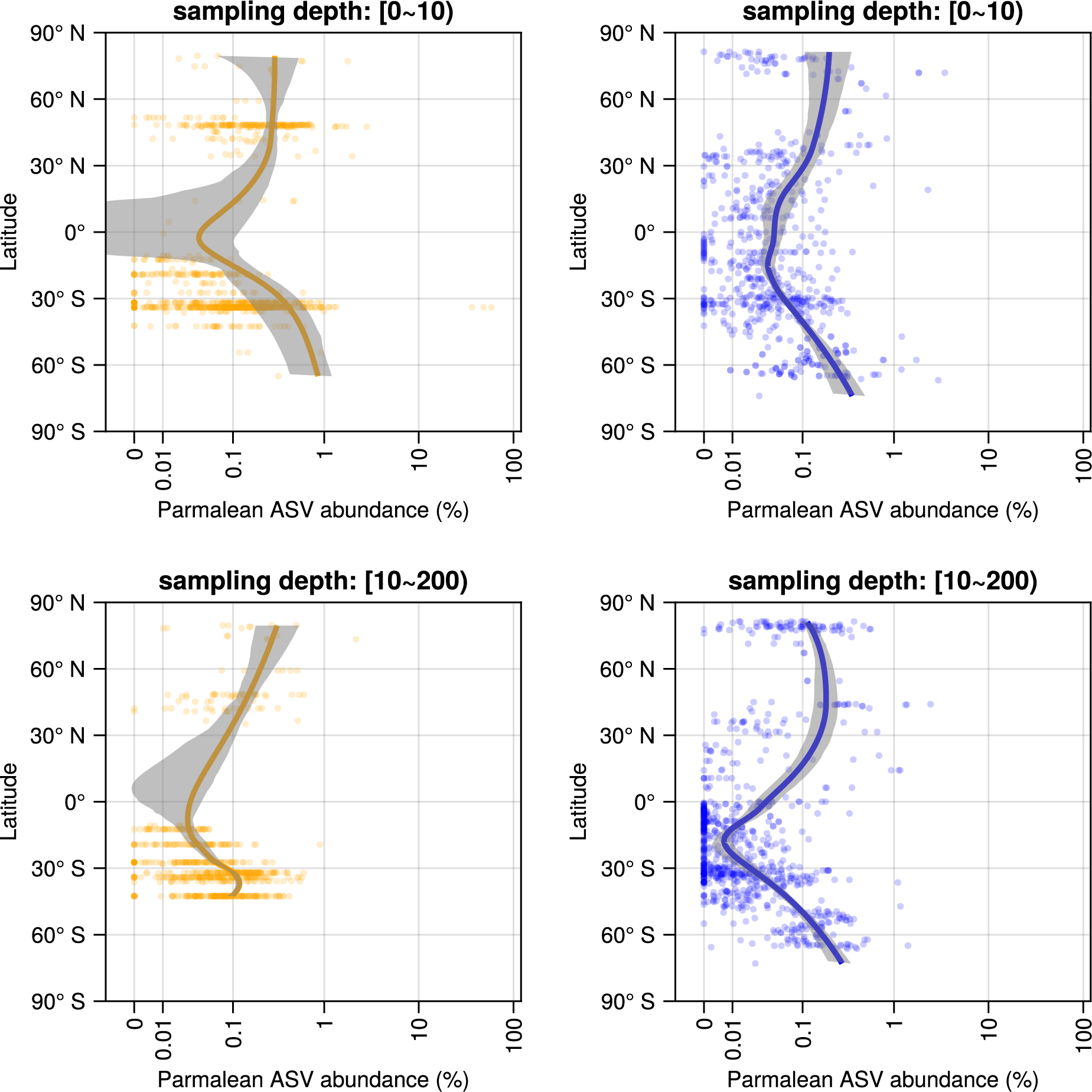
Latitudinal trend of total relative abundance of parmalean ASVs. Legend as in Fig. 3a for panels. Shaded areas represent 90 % confidence intervals.

### Global diversity pattern of parmalean ASVs

All Parmales clades were distributed in both the coastal and open oceans, but distinct distribution patterns emerged for the clades (Fig. 5). Clade I (‘Scaly parma’) appeared to be biased toward subarctic to polar regions (Fig. 5a). The WAT index of Clade I was 6.64℃, suggesting a preference for lower temperatures (Fig. 6a, Fig. S3a). Clade II (environmental clade) was rare in the tropic, but was widely distributed through mid- and high-latitude areas (Fig. 5b). The WAT index of Clade II is 14.2℃, which is higher than that of Clade I (Fig 6a, Fig. S3b).

**Fig. 5.**
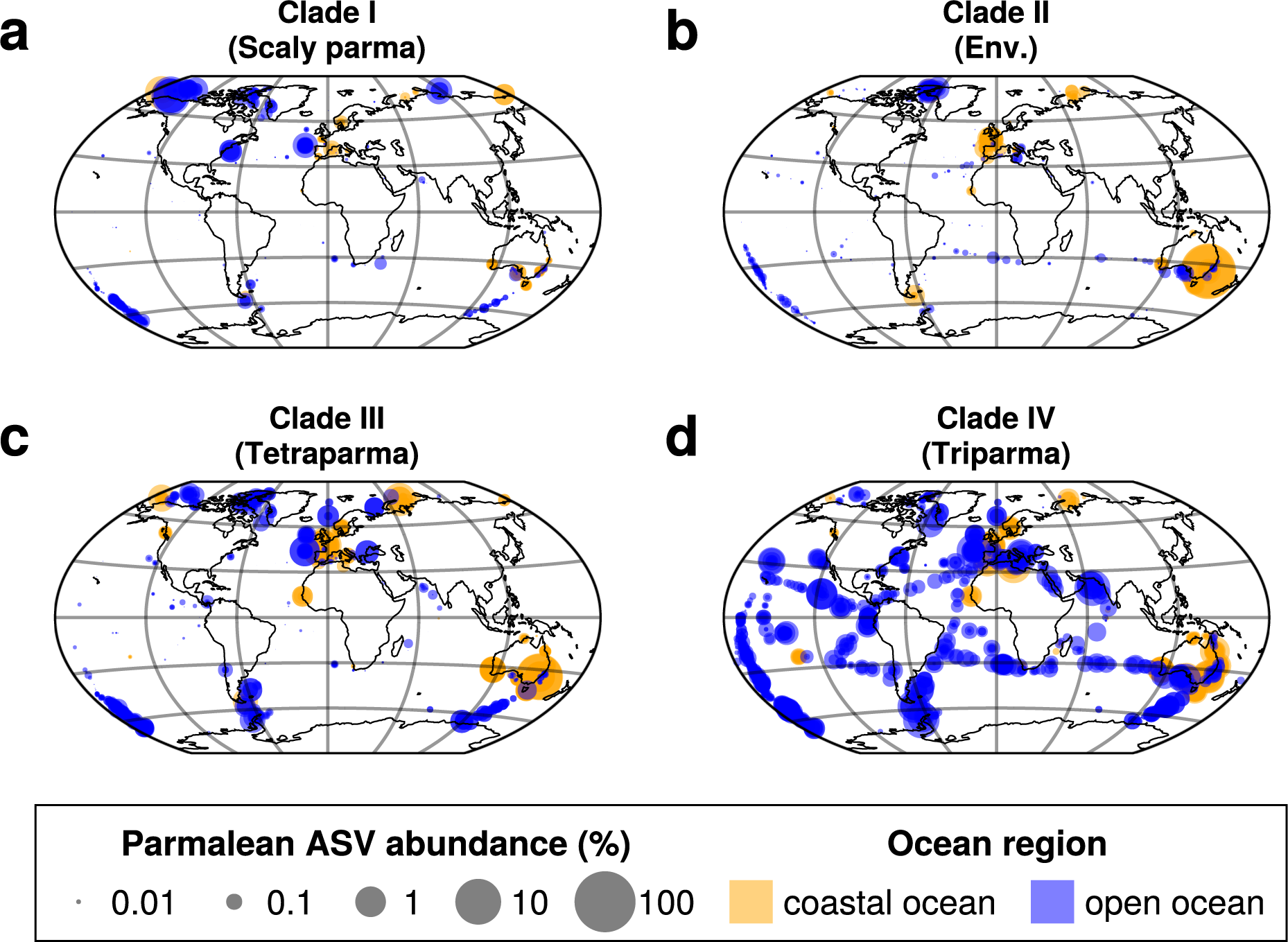
Global distribution of parmalean ASVs in each clade. Samples from all depths are show in single plots. Orange dots indicate coastal ocean samples and blue dots indicate open ocean samples. Marker sizes are scaled to the log10(total relative abundance*10,000+1) at each sample. (a) Clade I (‘Scaly Parma’), (b) Clade II (environmental clade), (c) Clade III (*Tetraparma*), (d) Clade IV (*Triparma*).

**Fig. 6.**
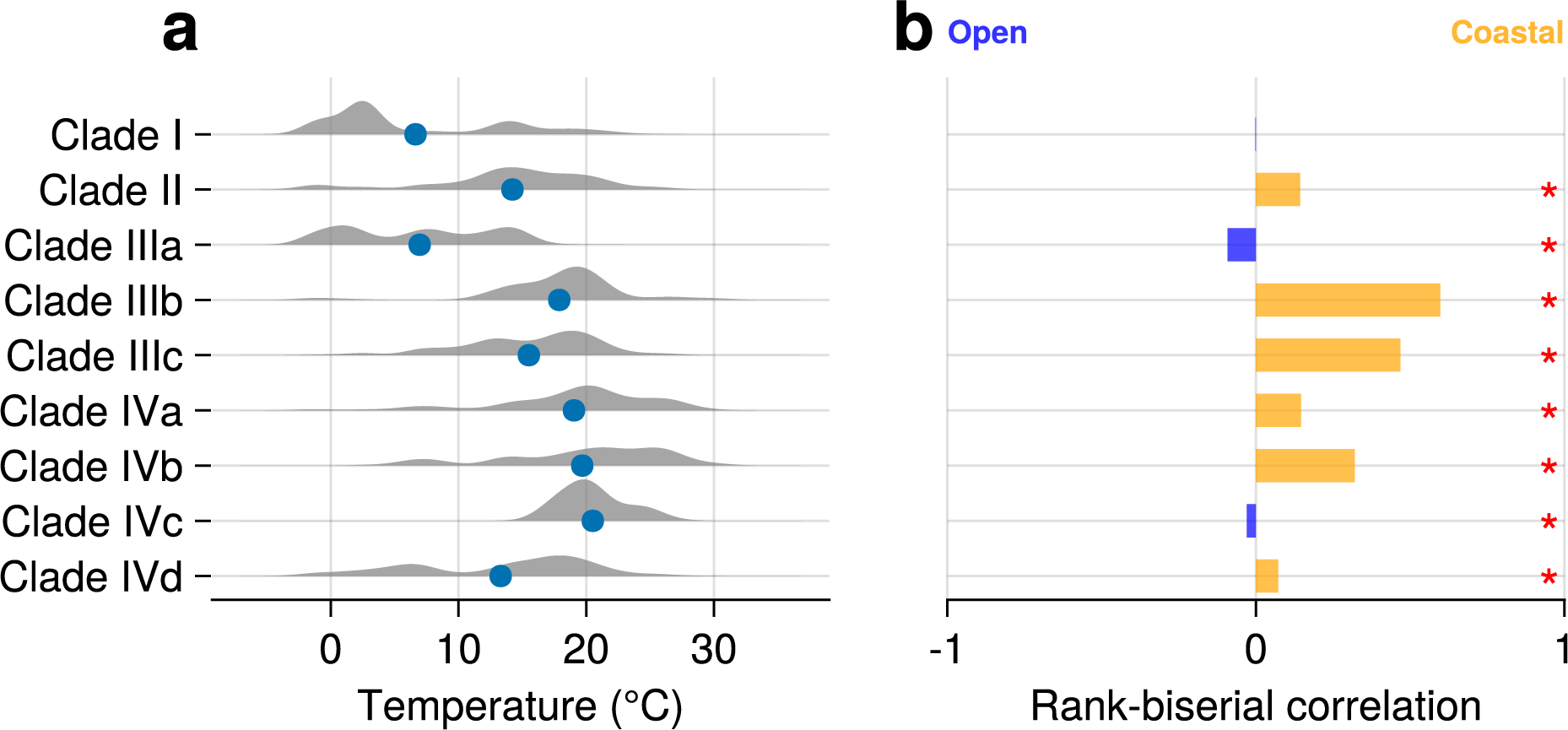
Characteristics of the distribution of clades/subclades. (a) Temperature preference of each clade. Blue dots represent the respective WAT indices. Shaded areas are violin plots of temperature weighted by the frequency in each sample. (b) Preference between coastal ocean and open ocean of each clade/subclade. Bar plot indicates rank-biserial correlation, which is an effect size of the Mann-Whitney U test. Positive RBC values indicate a preference for the coastal ocean, negative values indicate a preference for the open ocean, and the absolute value (0–1) indicates the strength of the preference. Asterisks indicate statistical significance of the test for each clade/subclade (*p*-value < 0.05). Detailed results of Mann-Whitney U test are in Table S2.

We conducted detailed subclade analysis of Clade III (*Tetraparma*) and Clade IV (*Triparma*) (Fig. 5c, d). Each of the three subclades of Clade III showed clearly distinct distribution patterns. Clade IIIa was distributed in both the coastal and open oceans from the subarctic to polar regions, suggesting a preference for cold water (Fig. 6a, 7a). Clade IIIb and Clade IIIc exhibited a strong bias towards coastal oceans (Fig. 7b, c), with the RBC values of Clade IIIb and Clade IIIc being 0.598 and 0.468, respectively (Fig. 6b, Table S2, Fig S5d, S5e).

**Fig. 7.**
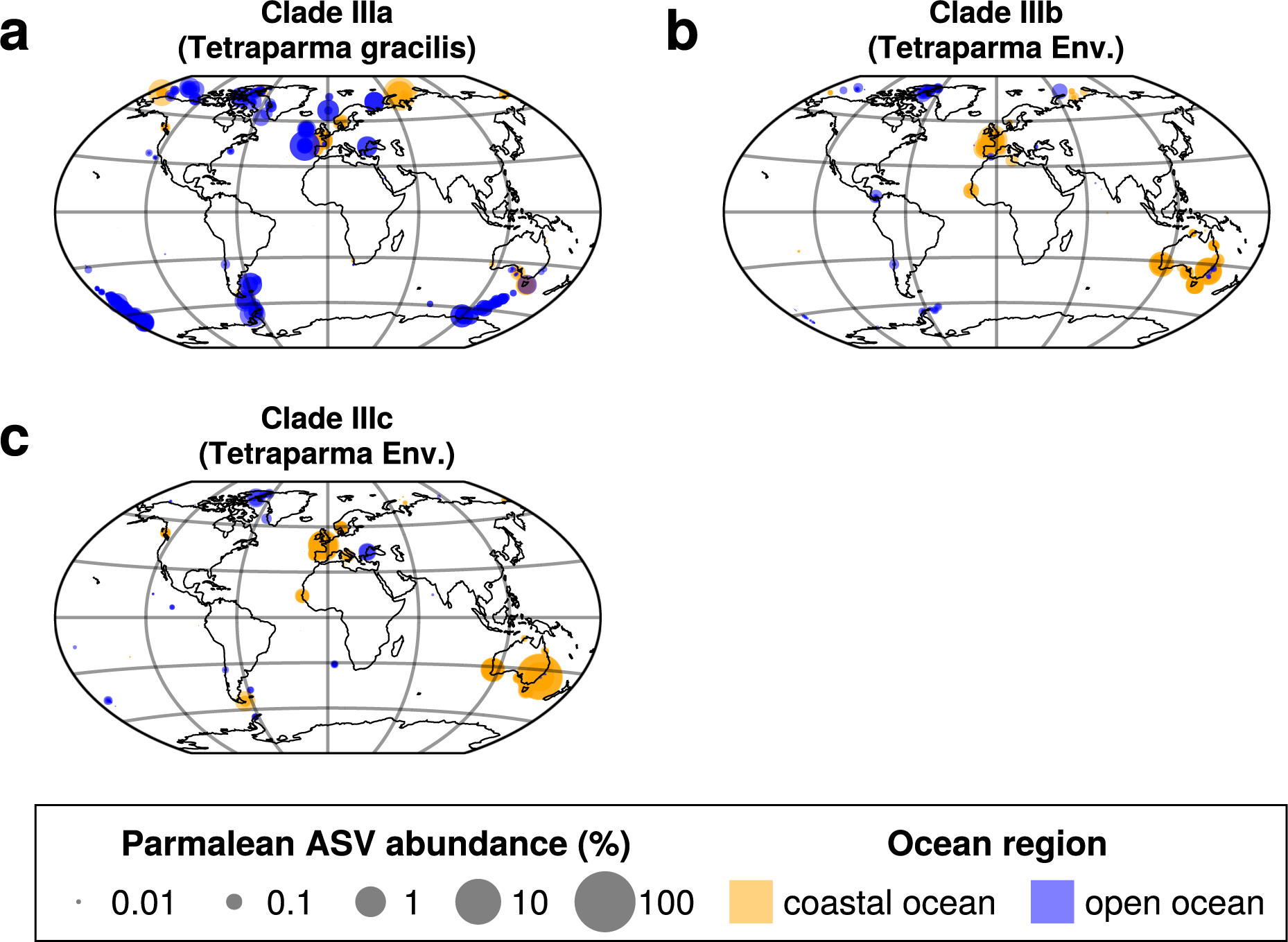
Global distribution of parmalean ASVs in each subclade of Clade III. Legend as in Fig. 5. (a) Clade IIIa (*Tetraparma gracilis*), (b) Clade IIIb (environmental clade), (c) Clade IIIc (*Tetraparma*).

Clade IVa, Clade IVb, and Clade IVd were very widely distributed, with each detected in over 50% of the samples (Fig. 8a, b, d). In contrast, Clade IVc was narrowly distributed and present in only 8.25% of samples. Clade IVc was only found in mid-latitude zones of both hemispheres, such as the Mediterranean Sea (Fig. 8c), and in a narrower range of water temperatures (Fig. 6a, Fig. S4f). Clade IVa and Clade IVb exhibited similar distribution patterns, being relatively rare in polar regions but still widely distributed (Fig. 8a, b); the WAT indices of the two clades were 19.0℃ and 19.7℃, respectively (Fig. 6a). The RBC values of Clades IVa and IVb were 0.146 and 0.320, respectively, with Clade IVb showing a stronger preference for the coastal oceans compared to Clade IVa (Fig. 6b, Table S2. Fig. S5f, S5g). Clade IVd was widely distributed but was less abundant in the tropics, in contrast to Clade IVa and Clade IVb. The WAT index of Clade IVb was 13.3℃, suggesting a preference for cold oceans (Fig. 6a, Fig. S4g).

**Fig. 8.**
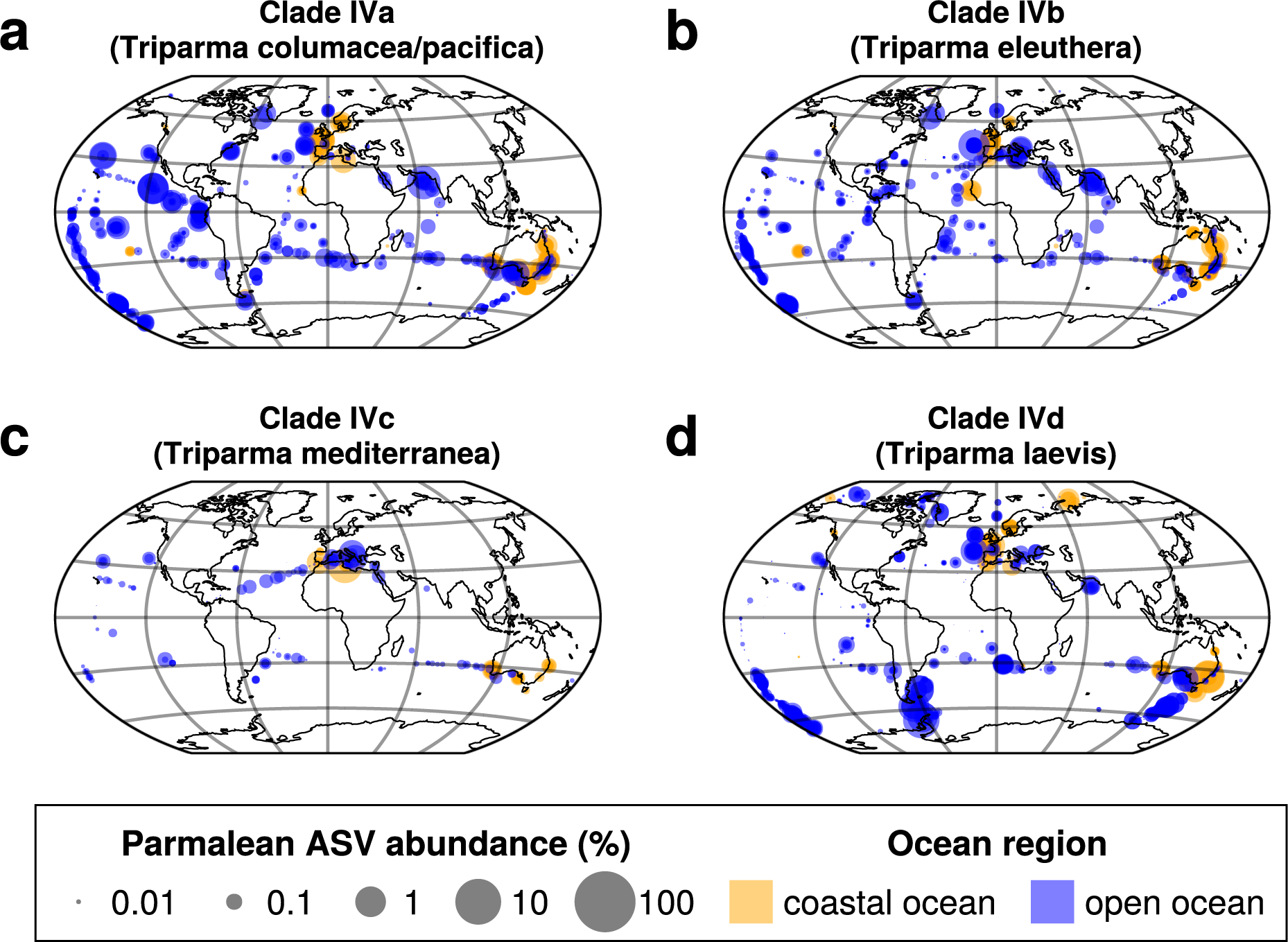
Global distribution of parmalean ASVs in each subclade of Clade IV. Legend as in Fig. 5. (a) Clade IVa (*Triparma columacea*, *Triparma pacifica*), (b) Clade IVb (*Triparma eleuthera*), (c) Clade IVc (*Triparma mediterranea*), (d) Clade IVd (*Triparma laevis*).

## Discussion

Using larger DNA metabarcoding datasets than those previously employed (Icnihomiya 2016; Kuwata 2018), we found that Parmales was broadly distributed from coastal to open oceans and from pole to pole (Fig. 3a). However, their relative abundances in the eukaryotic community were below 0.2% on average, as previously reported (Ichinomiya et al., 2016; Kuwata et al., 2018). Parmales can thus be considered a cosmopolitan phytoplankton group belonging to the rare biosphere (Lynch and Neufeld, 2015). Nonetheless, their relative abundance sometimes exceeds 1% (up to about 60%) (Fig. 3b), and they may occasionally play important ecological roles.

The predicted true richness of Parmales in the global ocean was about 316 ASVs. This low level of diversity contrasts starkly with their sister group, diatoms, which are estimated to have about 100,000 ribotypes of V9 region in the global ocean, methodological differences notwithstanding (Malviya et al., 2016). Therefore, Parmales is a minor group of eukaryotes with respect to diversity. The fitted Preston model suggests that the ASVs collected in this study covered 84.5% of the entire diversity of Parmales (Fig. 2b, c).

Previous field observations of silicified Parmales cells showed a strong association between the water temperature and their distribution (Ichinomiya and Kuwata, 2015; Ichinomiya et al., 2019). We revealed that the relative abundance of Parmales increases with latitude (Fig. 4), which supports the idea of an association between water temperature and Parmales distribution, although our analyses do not distinguish silicified forms from naked flagellates and thus differ from the previous reports in this regard.

The phylogenetic analysis of the full-length 18S rRNA gene sequences recovered four clades of Parmales, as previously reported (Kuwata et al., 2018; Ban et al., 2023) (Fig. 1). These sequences also include recent isolates of silicified forms (Ban et al., 2023), especially ‘Scaly parma’ and *Tetraparma gracilis*. Together with a previous analysis (Ban et al., 2023), our study confirmed the phylogenetic positions of ‘Scaly parma’ in Clade I and *Tetraparma* in Clade III (Fig. 1). Isolates of *Triparma* belonged to Clade IV. Clade II consisted only of environmental sequences with moderate levels of relative abundance (Fig. 2a) and a relatively high level of diversity (Fig. 2d, Table 1). *Pentalamina* (Booth and Marchant, 1987) is a genus of Parmales with a distinct morphology, but there have been no isolates or 18S rRNA sequences from this genus. Given the morphological differences in silicified cells among Clades I, III, and IV, *Pentalamina* may be a member of the environmental Clade II.

Individual clades/subclades showed distinct distribution patterns (Fig. 5, 6, 7, 8, Table S2, Fig S3, S4, S5), suggesting they were adapted to different ecological niches. Clade I (‘Scaly parma’) showed a strong preference for high-latitude regions and lower temperatures (Fig. 5a, 6a), which is consistent with previous findings (Kuwata et al., 2018) and our observation that the lone ‘Scaly Parma’ isolate derived from cold waters of the Sea of Okhotsk (Ban et al., 2023). Clade I is the least abundant among the four clades (Fig. 2a). The lack of field observations (apart from the single isolate) may be explained by this low abundance.

Clade II (environmental clade) also showed a preference for cold water, though not to the extent of Clade I (Fig. 5b, 6a). In samples where Parmales dominated in Botany Bay, Sydney, Australia, the most frequent ASV belonged to Clade II. This suggests Clade II members have the ability to cause bloom. As mentioned above, we speculate that *Pentalamina* belongs to Clade II. The silicified form of *Pentalamina* has only been reported from the Antarctic Ocean (Kuwata et al., 2018), which is consistent with the niche preference of Clade II.

Clade III (*Tetraparma*) and Clade IV (*Triparma*) showed distinct distribution patterns when divided into subclades (Fig. 6, 7, 8). Clade IIIa (*Tetraparma gracilis*) preferred cold water (Fig. 6a, 7a), which is consistent with reports that the silicified form of *Tetraparma gracilis* was frequently observed in cold water surrounding Hokkaido, North Japan (Ichinomiya et al., 2019). Clade IIIb and Clade IIIc showed a different distribution pattern; they were preferentially distributed in coastal areas (Fig. 6b, 7b, 7c, Table S2, Fig S5f, S5g).

Clade IVc (*Triparma mediterranea*) prefers the mid-latitude of both hemispheres (Fig. 8c). It was previously proposed that the distribution of *Triparma mediterranea* was mostly restricted to the Mediterranean Sea (Ichinomiya et al., 2016; Kuwata et al., 2018). Our study reveals that the mid-latitude distribution is a feature of Clade IVc, suggesting that parmaleans of this clade are restricted to a narrow temperature range (Fig. 6a). Clades IVa, IVb, and IVd were widely distributed and four of the highest sample coverage ASVs were from these clades (Fig. 3c).

Clade IVd (*Triparma laevis*) showed a preference for cold water, as previously reported (Ichinomiya et al., 2016; Kuwata et al., 2018), while Clade IVa (*Triparma columacea*, *Triparma pacifica*) and Clade IVb (*Triparma eleuthera*) showed a preference for warmer water. This result is consistent with previous growth experiments; silicified form isolates of *Triparma laevis* f. *inornata*, *Triparma* laevis f. *longispina*, and *Tripamra strigata* (all in Clade IVd) can grow in cold water but not over 15℃ (Ichinomiya and Kuwata, 2015), while the naked flagellate isolate of *Triparma eleuthera* (Clade IVb) can grow at 16–24℃ (Stawiarski et al., 2016). Previous observations of *Triparma columacea* and *Triparma retinervis* (each in Clade IVa) in the tropics (Fujita and Jordan, 2017) are also consistent with our results on the distribution of Clade IV. With the exception of Clade IIIb and Clade IIIc (coastal groups), all clades/subclades commonly appeared in both coastal and open oceans (Fig. 5, 6b, 7, 8, Table S2, Fig. S5), suggesting that many parmaleans are able to adapt to both eutrophic coastal oceans and nutrient-depleted open ocean. Parmales may switch between silicified photoautotrophic and naked flagellated phago-mixotrophic stages in their life cycle (Ichinomiya et al., 2016; Ban et al., 2023), and mixotrophs are generally thought to widen their niche by alternating their trophic strategies (Endo et al., 2018; Xu et al., 2022). Therefore, our results corroborate the idea that the cosmopolitan distribution of parmaleans might be explained by their life cycle strategy (Ban et al., 2023).

By integrating the large dataset produced by EukBank with prior morphological and genetic information, we firmly established that Parmales is a cosmopolitan but rare group of microeukaryotes that can occasionally make blooms. The mapping of morphological features onto the phylogenetic tree revealed still sparse but consistent signals supporting the correspondence between the clades and the different morphologies. Different clades display distinct spatial distributions, suggesting niche differentiations during the evolution of Parmales. We believe that the biogeography of different clades of Parmales revealed in this study will inform our understanding of the physiology, ecology, and evolution of Parmales.

**The EukBank Team Members and Affiliations**

Cédric Berney^1^, Frédéric Mahé^2,3^, Nicolas Henry^4,5^, Colomban de Vargas^6,5^

^1^Sorbonne Université, CNRS, Station Biologique de Roscoff, AD2M, UMR 7144, ECOMAP, Roscoff, France

^2^CIRAD, UMR PHIM, F-34398, Montpellier, France

^3^PHIM, Univ Montpellier, CIRAD, INRAE, Institut Agro, IRD, Montpellier, France.

^4^CNRS, Sorbonne Université, FR2424, ABiMS, Station Biologique de Roscoff, 29680 Roscoff, France

^5^Research Federation for the study of Global Ocean Systems Ecology and Evolution, FR2022/Tara Oceans GOSEE, 3 rue Michel-Ange, 75016 Paris, France

^6^Sorbonne Université, CNRS, Station Biologique de Roscoff, AD2M, UMR 7144, 29680 Roscoff, France

## Acknowledgments

We thank the *Tara* Oceans consortium, the EukBank consortium, and the people and sponsors who supported the Tara Oceans Expedition (http://www.embl.de/tara-oceans/) for making the data accessible. This is contribution number XXX of the *Tara* Oceans Expedition 2009–2013. This work was supported by the JST “Establishment of University Fellowships Towards The Creation of Science Technology Innovation” Grant Number JPMJFS2123, JSPS KAKENHI (No. 17H03724, 21K12231), and the Collaborative Research Program of Institute for Chemical Research, Kyoto University (Nos. 2015-39, 2016-30, 2020-33). Computational time was provided by the SuperComputer System, Institute for Chemical Research, Kyoto University. We are grateful to the Roscoff Bioinformatics platform ABiMS (http://abims.sb-roscoff.fr), part of the Institut Français de Bioinformatique (ANR-11-INBS-0013) and BioGenouest network, for providing computing and storage resources. We thank J. Carlson, PhD, from Edanz (https://jp.edanz.com/ac) for editing a draft of this manuscript.

## Supplementary information

## Supplementary Note

### Depth data preprocessing

In this study, in order to maximize the number of samples for analysis, depth categories in metadata and information from the original papers were used for samples without sampling depth values. The information was used to classify the samples into two categories: 0 m to less than 10 m (surface layer) and 10 m to less than 200 m (euphotic zone). Specifically, the categorizing proceeded as follows. Initially, all samples with depth values were systematically categorized to their respective depths. Then we eliminated samples that lacked both depth and depth category data. For the samples without depth data but with the depth category data being ‘[SRF] surface water layer (ENVO_00010504)’, only samples under project pohem (Ramond et al., 2019, 2021) were retained and categorized to the surface layer, because the original papers documented that these samples were collected at 0–5 m. Other samples were removed because they either deviated from the specified range, lacked the necessary descriptions in the original papers, or lacked original papers. For the samples without depth data but with the depth category data being ‘[DCM] deep chlorophyll maximum layer (ENVO_01000326)’, all samples are retained and categorized to the euphotic zone. For the samples without depth data but in other depth categories, all samples were removed.

## Supplementary Figures

**Fig. S1.**
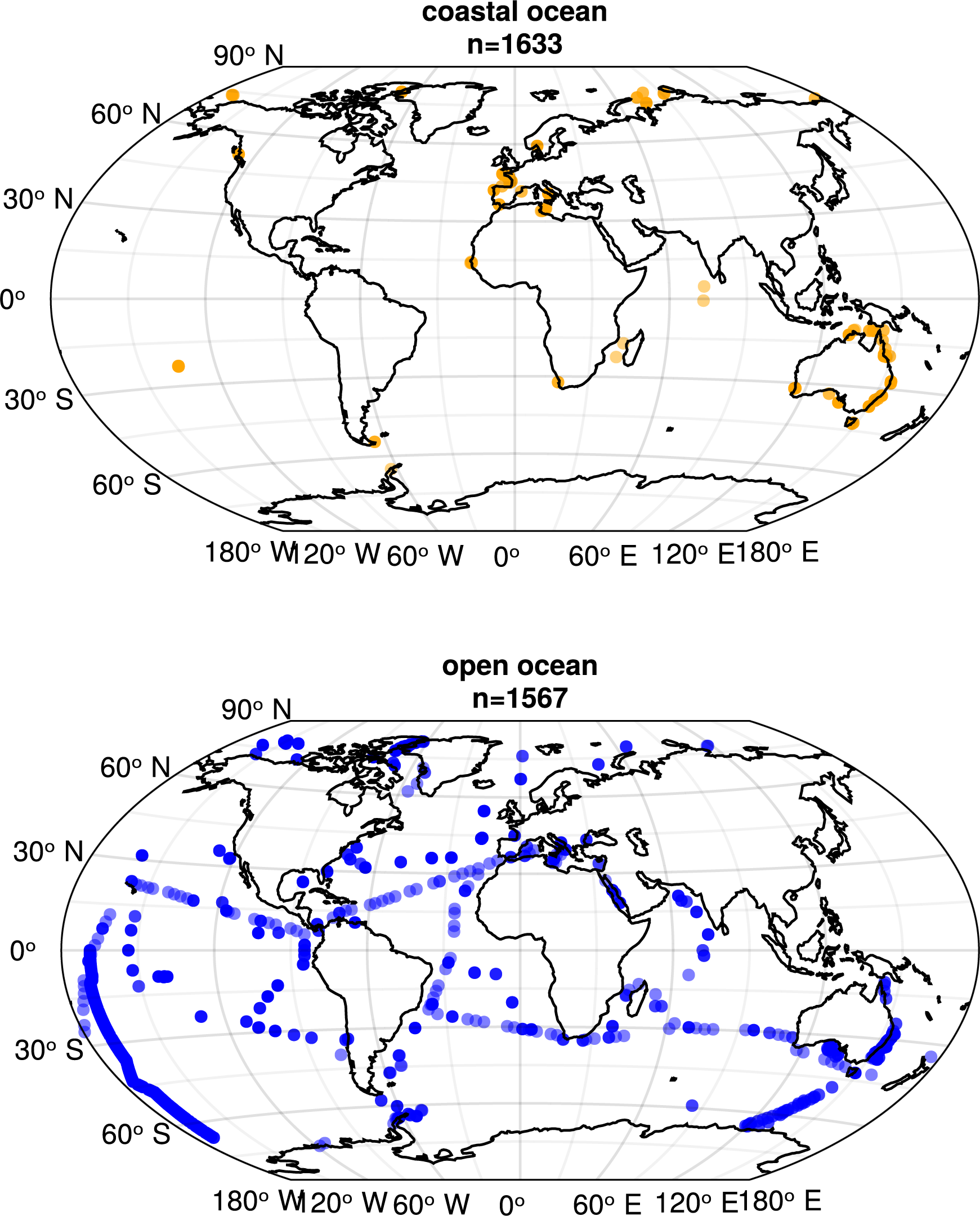
Geographic distribution of ocean samples selected from EukBank. Orange dots represent coastal ocean samples and blue dots represent open ocean samples.

**Fig. S2.**
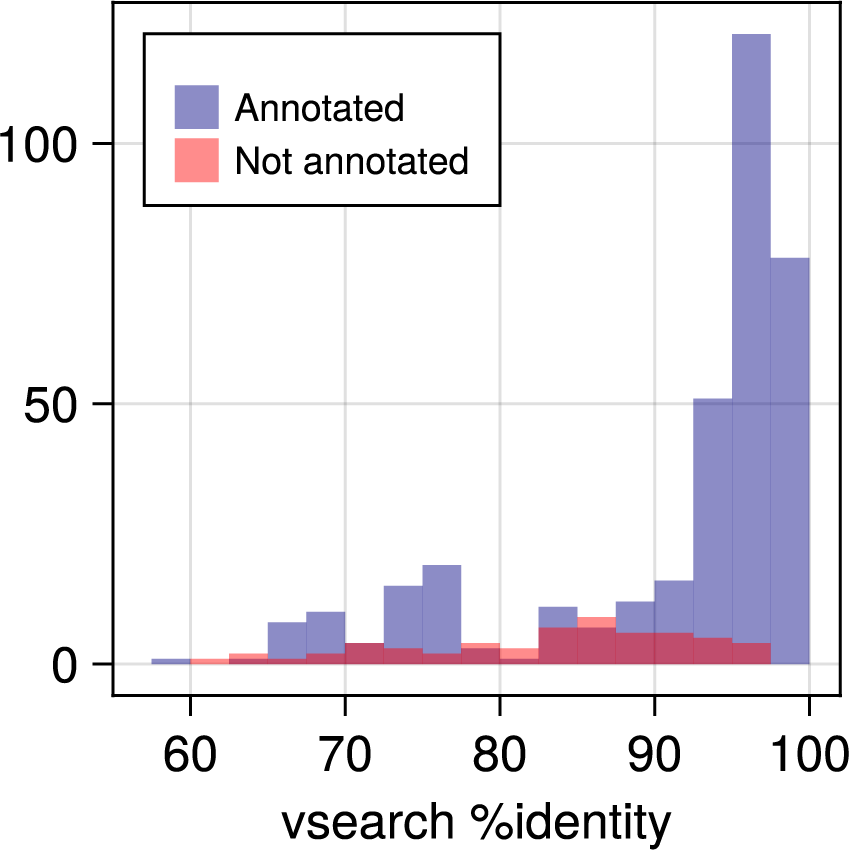
Distribution of identity against the best hit. Histogram illustrating the distribution of identity (%) against the best hits using vsearch. Blue bars represent ASVs annotated to the four clades, while red bars represent unannotated ASVs.

**Fig. S3.**
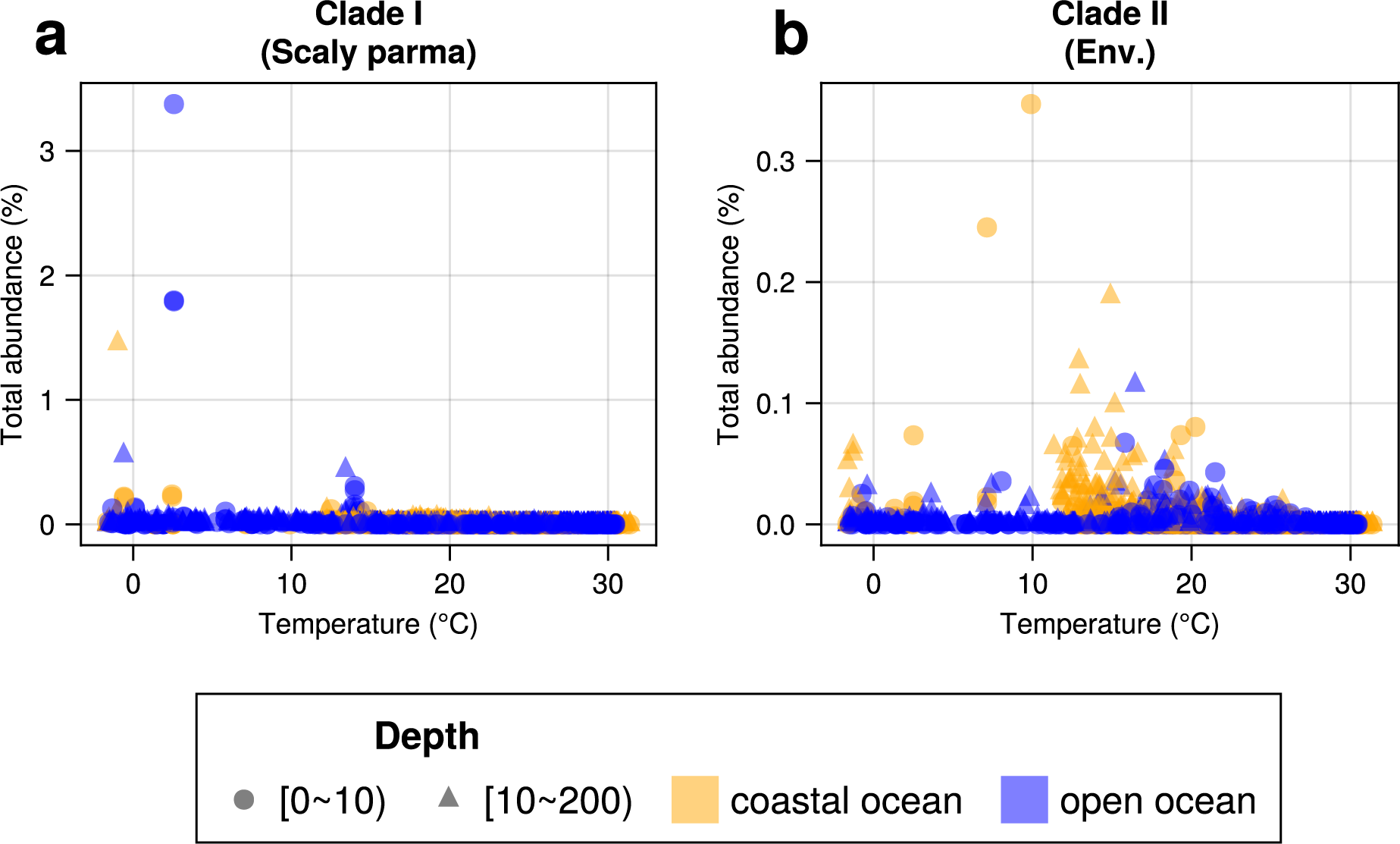
Distribution of total abundance across water temperature for Clade I and Clade II. Circles represent the surface (0-10 m), and triangles represent the euphotic zone (10-200m). Orange markers indicate coastal ocean samples and blue markers indicate open ocean samples. (a) Clade I (‘Scaly Parma’), (b) Clade II (environmental clade).

**Fig. S4.**
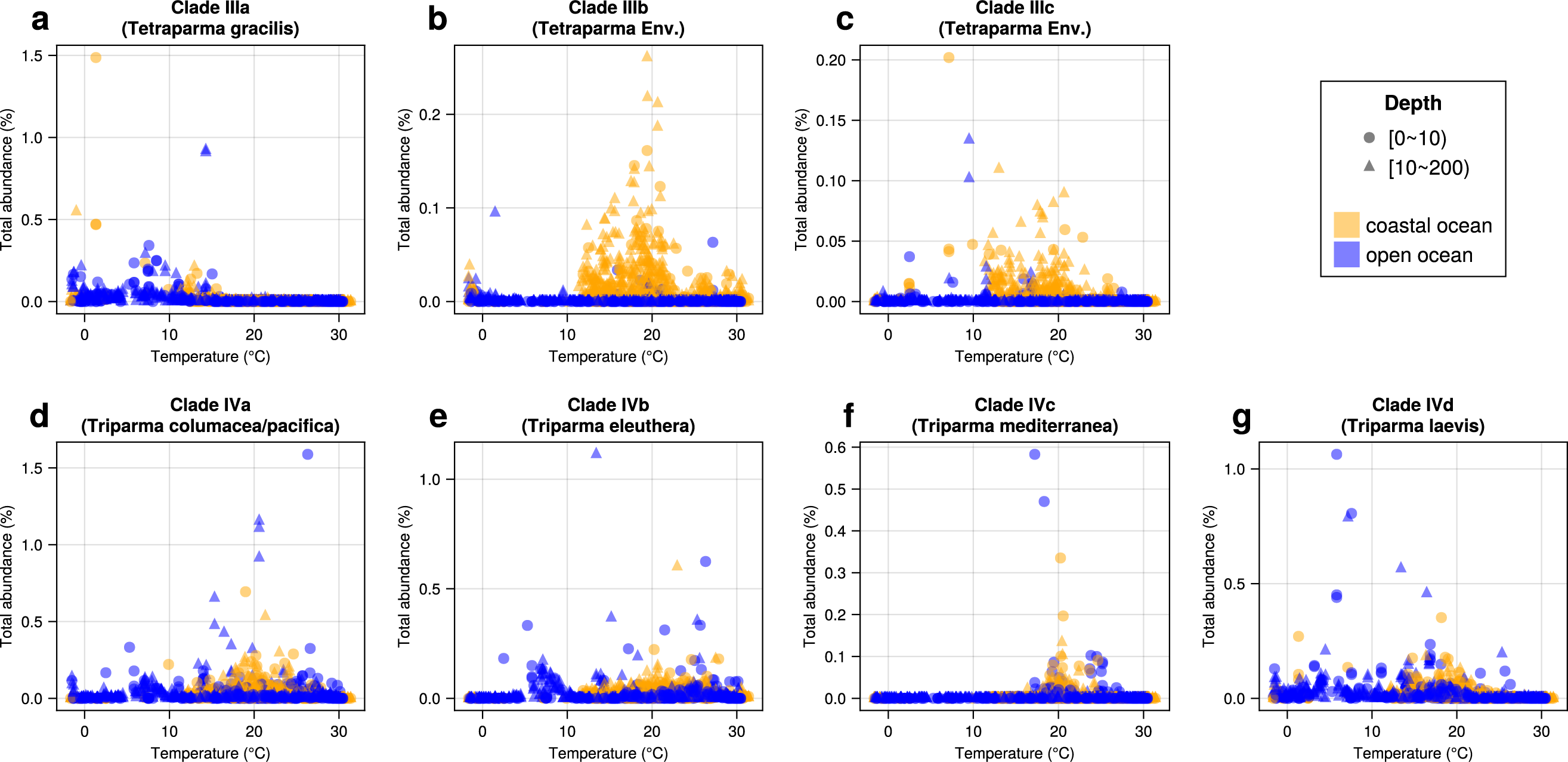
Distribution of total abundance across water temperature for each subclade of Clade III and Clade IV. Legend as Fig S3. (a) Clade IIIa (*Tetraparma gracilis*), (b) Clade IIIb (environmental clade), (c) Clade IIIc (environmental clade), (d) Clade IVa (*Triparma columacea*, *Triparma pacifica*), (e) Clade IVb (*Triparma eleuthera*), (f) Clade IVc (*Triparma mediterranea*), (g) Clade IVd (*Triparma laevis*).

**Fig. S5.**
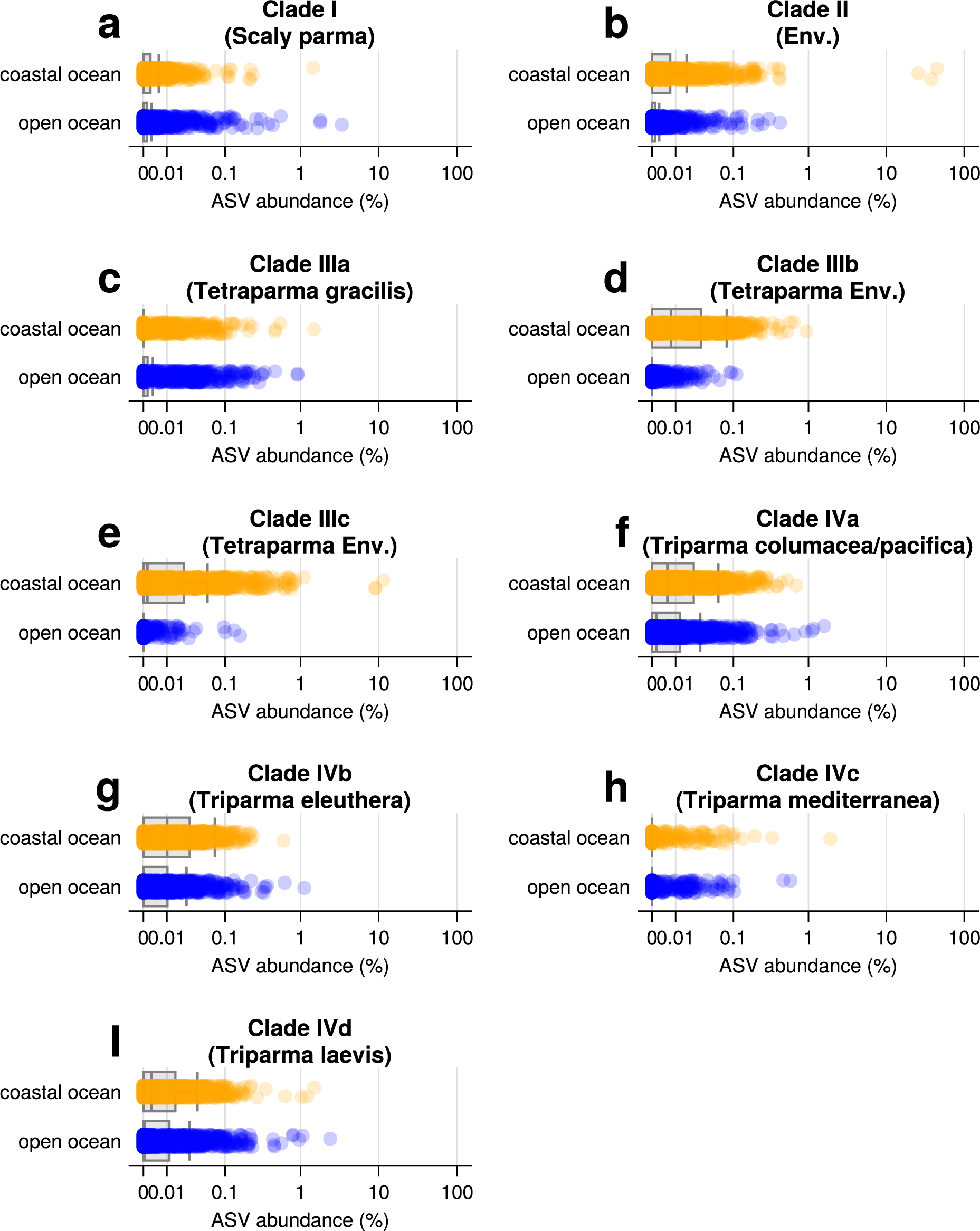
The distribution of total abundance in the coastal ocean and open ocean of each clade/subclade. (a) Clade I (‘Scaly Parma’), (b) Clade II (environmental clade), (c) Clade IIIa (*Tetraparma gracilis*), (d) Clade IIIb (environmental clade), (e) Clade IIIc (environmental clade), (f) Clade IVa (*Triparma columacea*, *Triparma pacifica*) (g) Clade IVb (*Triparma eleuthera*), (h) Clade IVc (*Triparma mediterranea*), (I) Clade IVd (*Triparma laevis*).

**Table S1.**
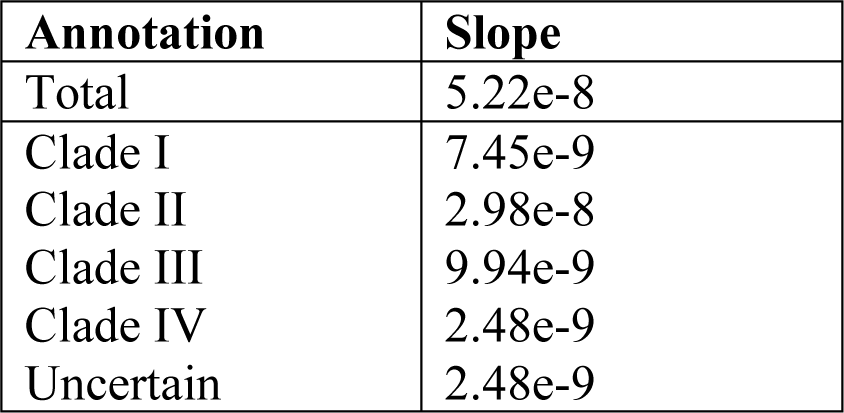
Slopes of each rarefaction curve.

| **Annotation** | **Slope** |
| --- | --- |
| Total | 5.22e-8 |
| Clade I | 7.45e-9 |
| Clade II | 2.98e-8 |
| Clade III | 9.94e-9 |
| Clade IV | 2.48e-9 |
| Uncertain | 2.48e-9 |

**Table S2.**
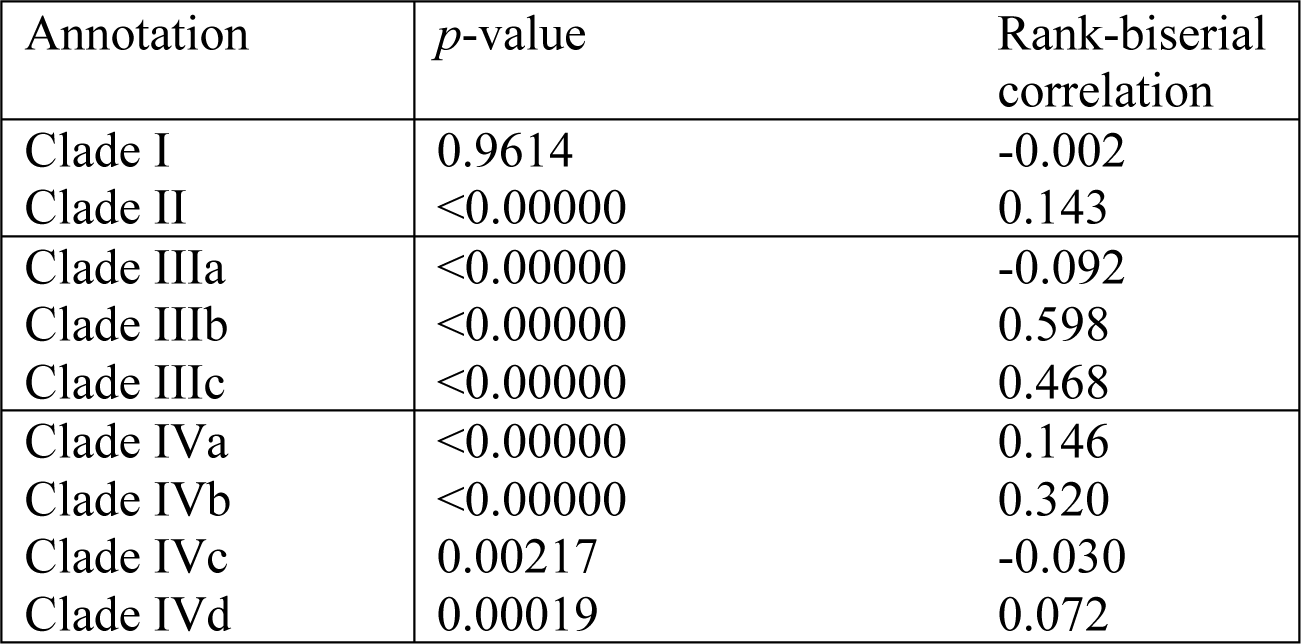
Results of Mann-Whitney U test comparing relative abundance between coastal ocean and open ocean.

| Annotation | *p*-value | Rank-biserial correlation |
| --- | --- | --- |
| Clade I | 0.9614 | -0.002 |
| Clade II | <0.00000 | 0.143 |
| Clade IIIa | <0.00000 | -0.092 |
| Clade IIIb | <0.00000 | 0.598 |
| Clade IIIc | <0.00000 | 0.468 |
| Clade IVa | <0.00000 | 0.146 |
| Clade IVb | <0.00000 | 0.320 |
| Clade IVc | 0.00217 | -0.030 |
| Clade IVd | 0.00019 | 0.072 |

